# Temperature drives the evolution and global distribution of avian eggshell colour

**DOI:** 10.1101/559435

**Authors:** Phillip A. Wisocki, Patrick Kennelly, Indira Rojas Rivera, Phillip Cassey, Daniel Hanley

**Author notes:** **Correspondence and requests for materials** should be addressed to D.H.

## Abstract

The survival of a bird’s egg depends upon its ability to stay within strict thermal limits. Avian eggshell colours have long been considered a phenotype that can help them stay within these thermal limits^1,2^, with dark eggs absorbing heat more rapidly than bright eggs. Although disputed^3,4^, evidence suggests that darker eggs do increase in temperature more rapidly than lighter eggs, explaining why dark eggs are often considered as a cost to trade-off against crypsis^5–7^. Although studies have considered whether eggshell colours *can* confer an adaptive benefit^4,6^, no study has demonstrated evidence that eggshell colours *have* actually adapted for this function. This would require data spanning a wide phylogenetic diversity of birds and a global spatial scale. Here we show evidence that darker and browner eggs have indeed evolved in cold climes, and that the thermoregulatory advantage for avian eggs is a stronger selective pressure in cold climates. Temperature alone predicted more than 80% of the global variation in eggshell colour and luminance. These patterns were directly related to avian nesting strategy, such that all relationships were stronger when eggs were exposed to incident solar radiation. Our data provide strong evidence that sunlight and nesting strategies are important selection pressures driving egg pigment evolution through their role in thermoregulation. Moreover, our study advances understanding of how traits have adapted to local temperatures, which is essential if we are to understand how organisms will be impacted by global climate change.

---

The impact of global climate patterns on the evolution and distribution of traits is an area of increasing importance as global temperatures continue to rise. Birds’ eggs are an ideal system for exploring the intersection between climate and trait diversity, because a tight thermal range is necessary for the survival of the developing embryo^8^, as eggs are unable to regulate their own temperature^9^. As a result, many birds have adapted incubation behaviours and nest characteristics in response to local conditions^10,11^. In addition to these behavioural adaptations, the adaptive value of eggshell coloration for thermoregulation has been of longstanding interest^1,2,5^. These eggshell colours are generated by just two pigments^12^ and eggshell coloration is known to reflect local environmental conditions^13^.

The white colour found on many eggs (e.g., ostrich eggs) reflects incident solar radiation from their surfaces, but can draw the attention of predators^14^. By contrast, dark brown or heavily speckled eggs (e.g., artic loon eggs) may escape the visual detection of predators, particularly in ground nesting birds^15^, but these darker eggs should heat more rapidly when left in the sun^1,16^. Therefore, in hot climes the thermal costs must be balanced against the adaptive benefits of cryptic pigmentation, while in cold climes thermoregulation and crypsis provide synergistic benefits to birds laying dark brown eggs. Thus, the potential trade-off between thermal constraints and crypsis are not equivalent across the globe; eggs found near the poles should be darker, while those found near the equator should have higher luminance (appear brighter) and more variable colours. The strength of these relationships should covary with nest types, such that they are stronger in nests exposed to more light.

To examine these ecogeographic patterns, we quantified egg colours across their known geographic ranges. To accomplish this we generated coordinates of avian eggshell coloration within an opponent colour space spanning 634 species, representing 32 of the extant 36 orders of birds^17^ (Fig. 1). Coordinates within this space correspond with avian perceived colour and luminance (brightness), and they directly relate to physical metrics of colour (see Methods). Then, we simulated random nests (*n =* 3,577,243) within each species’ breeding range and assigned each species’ eggshell colour and luminance. Next, we calculated the phylogenetic mean colour and luminance (Fig. 1) for species found within each sampling area of an equal area hexagonal grid, and associated annual temperature, and other climatic variables, with these eggshell phenotypes. We then used a spatial Durbin error model to predict eggshell colours and luminance values by climate variables to account for spatial autocorrelation. Lastly, to explore the direct effects of solar heating on eggshell colours, we tested the heating and cooling rates of white, blue-green, and brown *Gallus gallus domesticus* eggs under natural sunlight conditions.

**Fig. 1.**
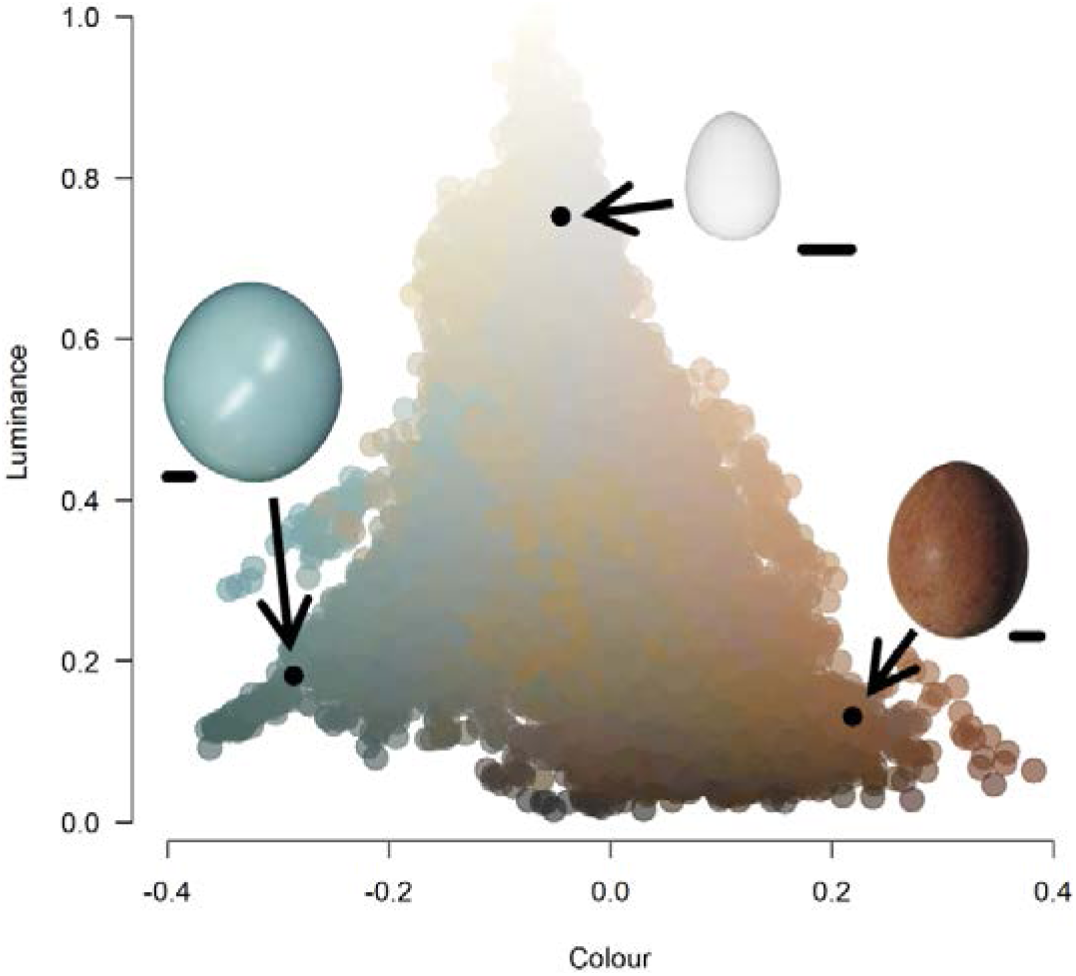
An opponent colour space illustrating the avian perceivable variation in eggshell coloration. These data are based on the difference between two opponent channels (colour) and avian perceivable luminance. Inset eggs represent where three distinct eggshell colour morphs fall within this space. We illustrate the locations for eggshell colours for the black tinamou *Tinamus osgoodi*, peregrine falcon *Falco peregrinus*, and Olive sparrow *Arremonops rufivirgatus*, representing blue-green, brown, and white egg colours, respectively. Each egg is depicted alongside a 1 cm scale bar.

We found that avian eggshell colours are darker and browner near the Arctic, and have greater luminance and more variable colours near the equator (Fig. 2). Temperature accounted for 83.3% and 88.0% of the variance in avian egg colour and luminance, respectively, with higher latitudes having significantly browner (*temperature*: z = −5.29, p < 0.0001; *temperature*^*2*^: z = 2.04, p = 0.04) and darker eggs (*temperature*: z = 13.32, p < 0.0001; *temperature*^*2*^: z = −8.50, p < 0.0001). These striking, nonlinear relationships with latitude and temperature suggest that avian eggshell colours are adaptive for thermal regulation in cold climes, but not in other environments. In support of this, we found direct linear associations between annual temperature and eggshell colour and brightness within two distinct climate regions that are associated with cold climates (*colour*: R^2^= 0.82, z = −9.30, p < 0.0001; *luminance*: R^2^ = 0.94, z = 14.79, p < 0.0001; Fig. 3a,b), while those patterns are weaker in other climate regions (*colour*: R^2^= 0.79, z = −3.03, p = 0.002; *luminance*: R^2^ = 0.72, z = 0.59, p = 0.56; Fig. 3c,d). Although the role of thermoregulation in driving egg colour evolution has long been proposed as an important selective pressure^1,2^, dark brown colours are often considered costly because egg temperatures are maintained close to their upper thermal limit^4^; thus, brown colours would be counterproductive to shedding incident heat. Instead, our results illustrate that this classic trade-off is dependent upon geography, where brown colours are adaptive for thermoregulation in only some places on Earth.

**Fig. 2.**
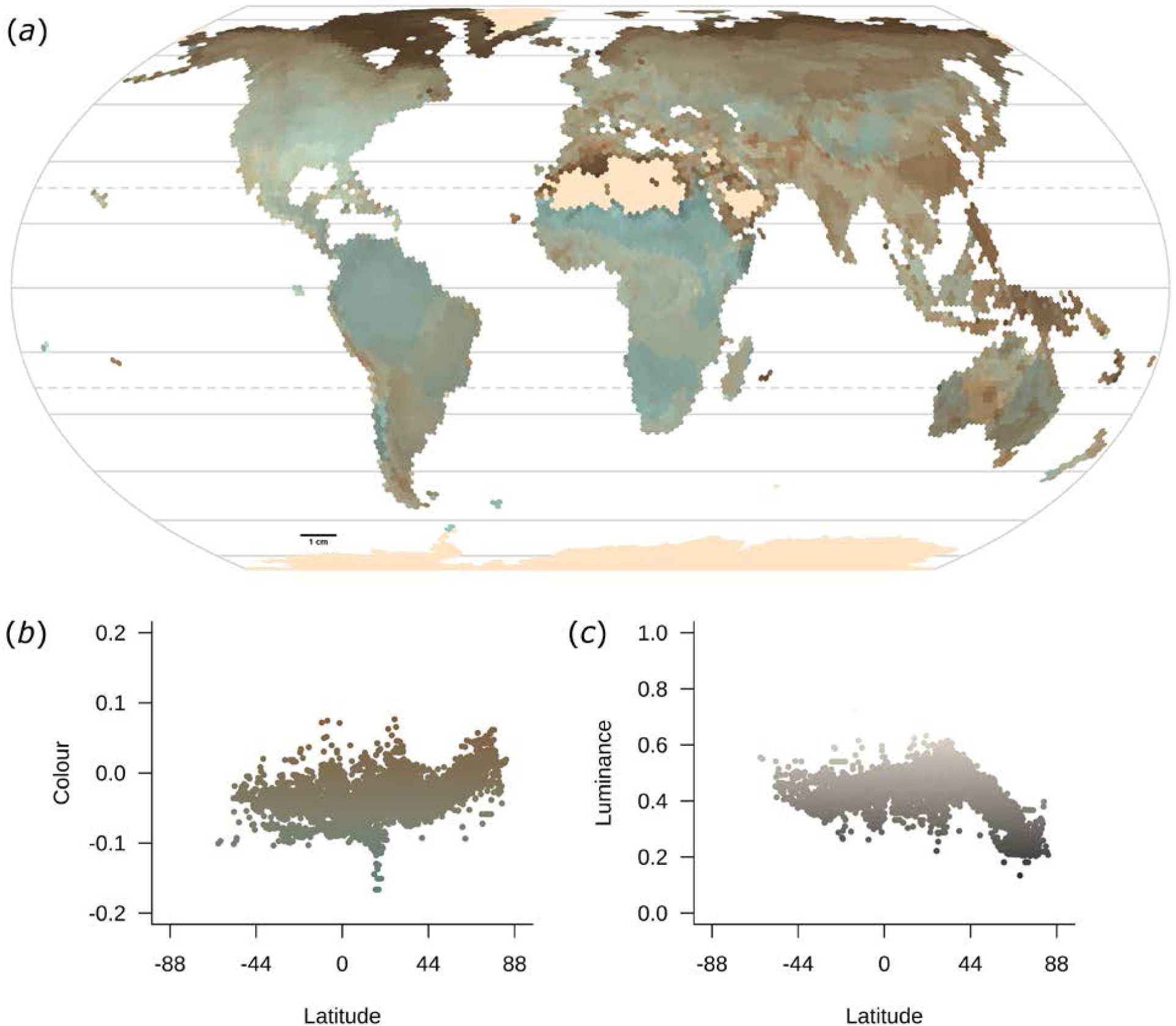
An equal Earth projection of the global distribution of avian eggshell colour. Depicted in a **a**, bivariate plot illustrating continuous variation in blue-green to brown eggshell and dark to light eggshell colours, in units of standard deviation from their means. Both avian perceived **b**, *colour* (R^2^= 0.83, *latitude*: z = −0.06, p = 0.95; *latitude*^*2*^: z = 4.67, p < 0.0001) and **c**, *luminance* (R^2^= 0.88, *latitude*: z = 2.08, p = 0.04; *latitude*^*2*^: z = −9.26, p < 0.0001) vary non-linearly across latitude, such that dark brown eggs are more likely at northerly latitudes.

**Fig. 3.**
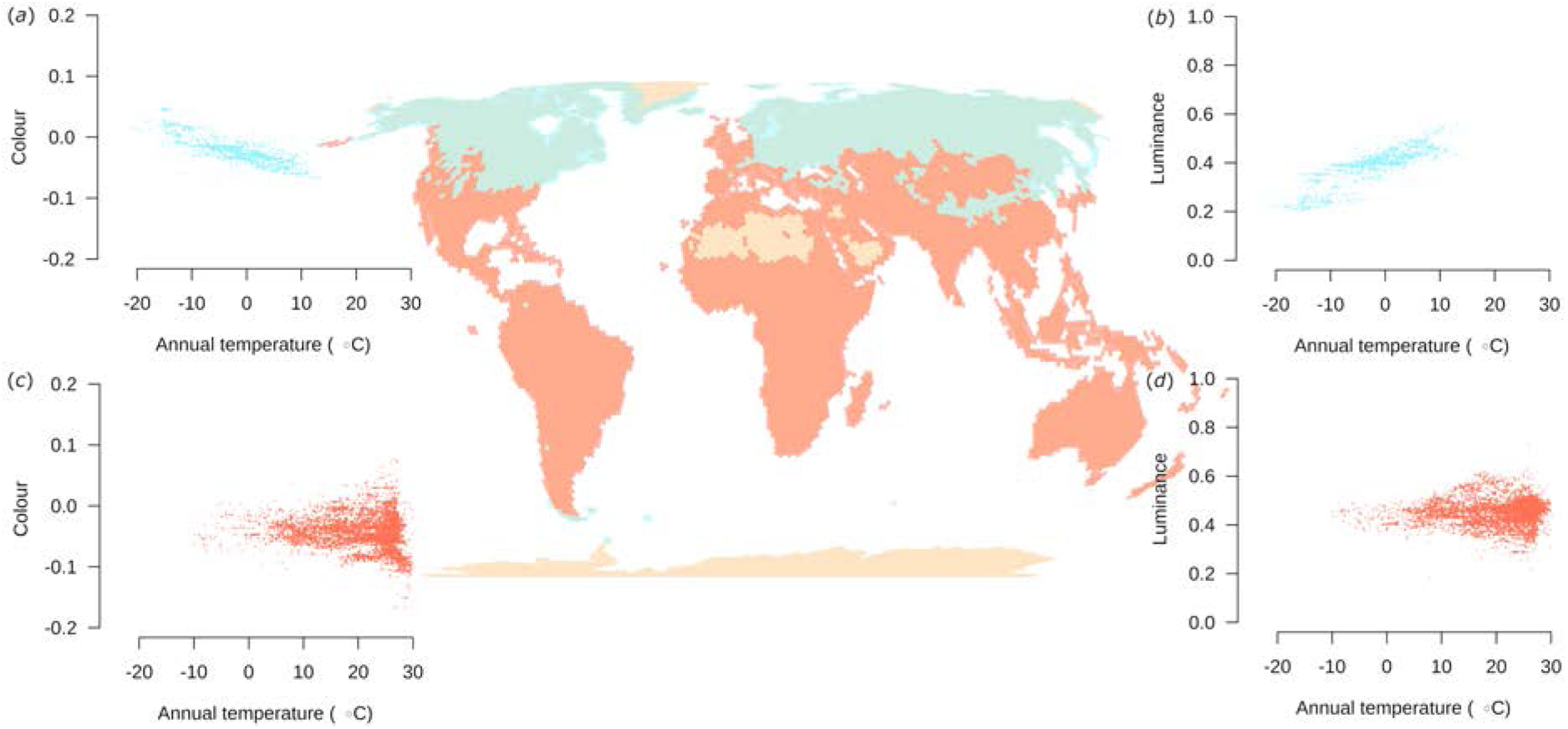
Global relationship between temperature and eggshell colour and luminance. Variation in avian perceived **a**,**c**, colour and **b**,**d**, luminance in **a**-**b**, cold Köppen climate regions (blue dots, *n =* 2,159) compared to **c**-**d**, other ecoregions (pink dots, *n =* 4,523). The central inset depicts those climate regions on the Earth. See Fig. 1 and methods for details.

Using a linear model, and under natural solar radiation, luminance and egg mass explained 91% of the variance in surface heat gain (F_2,45_ = 232.70, R^2^_adj_ = 0.91, p < 0.0001; *luminance*: z = −1.70, p < 0.0001; *mass*: z = 0.45, p = 0.003), while colour and mass only predicted 71% of the variance (F_2,45_ = 58.85, R^2^_adj_ = 0.71, p < 0.0001; *colour*: z = 1.15, p < 0.0001; *mass*: z = 0.53, p = 0.0005; ΔAICc = 55; Fig. 4). Specifically, dark brown eggs heated faster (7.28 °C h^−1^; F_3,44_ = 127.3, R^2^_adj_ = 0.89, p < 0.0001) than light brown (6.78 °C h^−1^; p = 0.006), blue-green (5.71 °C h^−1^; p < 0.0001), or white eggs (4.17 °C h^−1^; p < 0.0001) and retained heat longer (Extended Data Fig. 1a). This evidence suggests that pigmentation is an important factor in egg thermoregulation, such that darker eggs are more adaptive than lighter eggs in colder climes, especially in exposed nests. Because darker brown colours are more common in the coldest places, our findings suggest that the dark brown eggshell pigment, protoporphyrin, provides a greater thermal adaptive benefit than the blue-green eggshell pigment, biliverdin. Thus, birds may adapt an optimal colour to their locale.

**Fig. 4.**
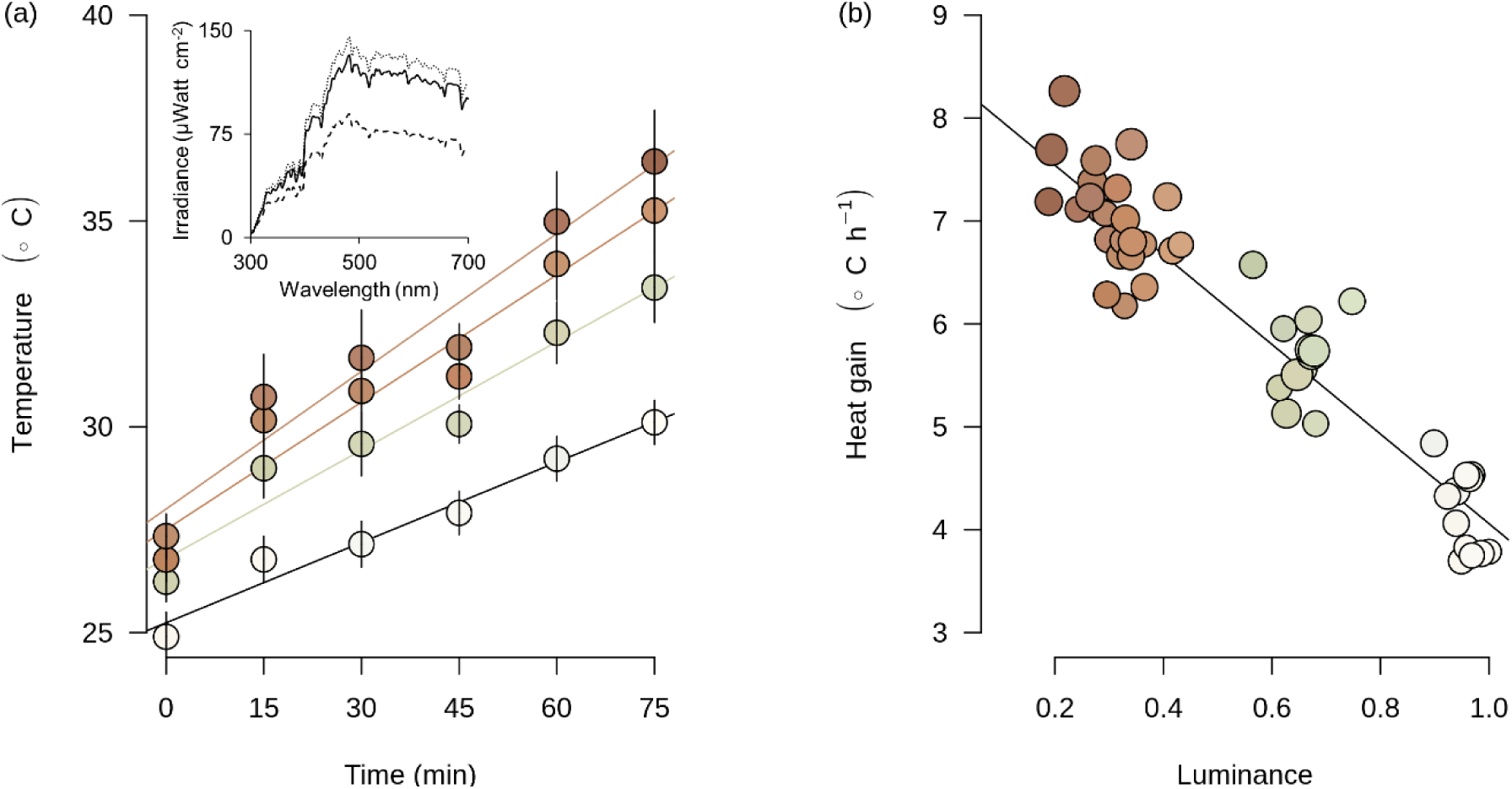
Eggshell heat gain. The heat gain ± s.e. for **a**, white, blue, light brown, and dark brown (bottom to top) chicken *Gallus gallus domesticus* eggs left outside at 27°C. The eggs were exposed to direct sunlight, except for cloud cover between 30-45 minutes after the start of the experiment (note the negative residuals at 45 minutes for each line). Inset includes solar irradiance measures at the start (solid black line), end (dotted grey line), and during cloudy conditions (dashed line) of the experiment. We also plot the **b**, eggshell surface heat gain in °C h^-1^ over this 75 min period, for each egg-based luminance and mass (dot size) illustrates relative egg sizes).

In cold climates, the ability to maintain temperature for longer periods of time afforded by darker coloration is particularly important^18^. This is not to say that species laying exposed eggs will leave their eggs unattended for longer, but instead, when unattended, dark eggs would have greater heat retention over comparable time periods. Eggshell pigmentation thus can confer an additional advantage over the chill tolerance found in some species^8^. By contrast, in warmer climates dark eggs might be more costly because they heat relatively quickly (e.g., nearly twice as fast as white eggs). In these environments, species are subjected to competing selection pressures and while eggs may have greater luminance (less pigmentation) in these warmer climes the colour is likely impacted by a range of other selective pressures: solar filtration^5,19^, anti-microbial defence^20^, signalling of mate quality^21^, and egg recognition^22^. Additionally, crypsis^7^and eggshell strength^23^ are known to influence egg coloration, and are likely important selective pressures globally. This interpretation is supported by our data. Egg colour was increasingly variable nearer the equator, indicating other selective pressures (e.g., ecological or behavioural) are acting on eggshell coloration.

As predicted, the strength of these relationships varied with nest types that experience differing levels of solar irradiance, such that the ability for temperature to predict colour and brightness was strongest in ground nesting birds which are exposed to the most light (*colour*: R^2^ = 0.77, z = −7.82, p < 0.0001; *brightness*: R^2^= 0.77, z = 9.63, p < 0.0001), was weaker in cup nesting birds that often nest in dense foliage (*colour*: R^2^= 0.62, z = 8.68, p < 0.0001; *brightness*: R^2^= 0.78, z = 10.93, p < 0.0001), and was weakest in cavity nests entirely enclosed from light (*colour*: R^2^ = 0.68, z = −1.02, p = 0.31, *brightness*: R^2^= 0.75; z = 2.46, p = 0.01). Interestingly, recent research has documented that in non-avian dinosaurs these two eggshell pigments emerged in species employing exposed nesting strategies^24^. This evidence suggests that nesting ecology was a pervasive and important selective pressure driving the evolution and distribution of eggshell colours.

The chemical properties of eggshell pigments underlying eggshell colours may have adapted in response to other environmental forces. For example, eggshell pigments may protect the developing embryo’s DNA from ionizing radiation, due to their absorption peaks in the UV range^5^. The blue-green pigment, biliverdin, absorbs more UV light than protoporphyrin (the brown pigment)^5^; therefore, we would expect more intense blue-green coloration in locales with high UVB radiation. Cold regions had on average 3.4 times less average monthly UVB radiation (mean UVB ± s.e.; *cold*: 276.54 ± 4.24 kJ·m^-2^; *other*: 943.82 ± 3.68 kJ·m^-2^) than other regions^25^, yet in both regions blue-green eggs were associated with locales with higher UVB and the relationship was strongest in cold climes which have lower UVB levels (*cold*: R^2^ = 0.82, z = −7.54, p < 0.0001; *other*: R^2^= 0.79, z = −4.28, p < 0.0001). Similarly, luminance was only related to UVB levels in cold climes (*cold*: R^2^ = 0.94, z = 11.24, p < 0.0001; *other*: R^2^ = 0.72, z = 0.44, p = 0.66), suggesting that these relationships are driven by the strong correlation between UVB and temperature (r = 0.91, CI_0.95_ = 0.91 to 0.92, p < 0.0001). Differentiating the independent selection pressures exerted by either is an important area of future research.

It is also possible that protoporphyrin protects the egg from microbial invasion because protoporphyrin has photo-dependent anti-microbial activity^20,26^. Because microbial loads are associated with humidity^27^, we expect to find browner eggs in more humid places. However, although we found that in cold regions humidity significantly predicted eggshell colour (R^2^ = 0.79, z = 3.80, p < 0.0002) and luminance (R^2^ = 0.93, z = −2.57, p = 0.01), these patterns were not as strong in other climate regions where the risk of microbial infection would be expected to be higher (*colour*: R^2^ = 0.79, z = 2.20, p = 0.03; *luminance*: R^2^= 0.72, z = 0.56, p = 0.58). Overall, our results indicate that temperature is the main selective pressure driving avian eggshell colour in colder northern climates, and that other selective pressures may be more important at warmer climates.

Here we provide a robust analysis across the full phylogenetic diversity of birds and at a global scale to consider how abiotic factors have shaped the evolution and distribution of a trait. We illustrate why such scale, scope, and depth is necessary to understand a classic example of an ecological trade-off. Thus, while our findings provide a framework for understanding the selective pressures shaping eggshell colours, they also provide insight into the forces driving pigmentation generally. We show that abiotic pressures such as temperature constrain the expression of phenotypes, and may limit the role of alternative selective pressures in some places while those same traits may be less constrained in other places on Earth. Such explorations of the impact of climate on phenotypes^28^, particularly those inherently linked with survival^29^, are necessary if we wish to quantify climate change impacts. As temperatures rise in the Arctic^30^, the egg colours found in that region could be shifting from adaptive to maladaptive, which could result in a loss of biodiversity. Therefore, our findings provide a roadmap for identifying regions at greater risk due to increasing global temperatures and prioritizing conservation efforts. Thus, in addition to illustrating how abiotic factors have shaped trait diversity, our research outlines novel and unexplored consequences of anthropogenic climate change.

## Online content

Any methods, additional references, Nature Research reporting summaries, source data, statements of data availability and associated accession codes are available at. Birdlife Range data can be requested at http://datazone.birdlife.org/species/requestdis. Digital Chart of the World basemap can be found at: https://worldmap.harvard.edu/data/geonode:Digital_Chart_of_the_World. Natural Earth Data maps can be found at: https://www.naturalearthdata.com/. Worldclim temperature data is available at http://worldclim.org/version2. National Center for Atmospheric Research UV data can be found at: https://www2.acom.ucar.edu/modeling/tuv-download. Atlas of the Biosphere humidity data can be found at: https://nelson.wisc.edu/sage/data-and-models/atlas/maps.php. Köppen climate data can be found at http://www.gloh2o.org/koppen/.

## Acknowledgements

We thank Long Island University for space and support for this project. We also thank K. Mendola, J. L. Cuthbert, and K. Dalto for their assistance, and S. Tettelbach, M. C. Stoddard, S. Madronich, and T.E. Lewis for helpful comments. We thank Mark Hauber (MEH) with assistance securing funding to collect eggshell spectra. The collection of the original dataset was partially funded by a Human Frontier Science Program (http://www.hfsp.org/) young investigators’ grant (RGY0069/2007-C) and a Leverhulme Trust (http://www.leverhulme.ac.uk/) project grant (F/00 094/AX) to PC and MEH.

## Author contributions

The concept was developed by D.H., data was collected by D.H. and P.C., colour analyses were conducted by D.H and I.R., biogeographic models were developed and implemented by P.W., P.K, and D.H., the initial draft was prepared by P.W. and D.H., and all authors edited the paper.

## Additional information

**Extended data** is available for this paper at _.

## METHODS

### Colour estimation

We used spectral reflectance data of avian eggshells (*n =* 634) spanning all avian orders excluding Eurypygiformes (two species: Kagu and Sunbittern), Leptosomiformes (one species: Cuckoo Roller), Mesitornithiformes (three Mesites species), and Pteroclidiformes (16 Sandgrouse species)^1^. Of those orders represented in our data, 87±4% of the families within them were sampled. These reflectance data were smoothed using a locally weighted polynomial function. These data had previously been used to determine the extant variation in avian eggshell coloration^17^ within the avian tetrahedral colour^32,33^. We then modelled avian perceived colour of each eggshell using a noise limited neural model to estimate quantum catch for each photoreceptor. We calculated relative photoreceptor and double cone quantum catch^34^ assuming the average photoreceptor sensitivity of an ultraviolet sensitive bird and the double cones of the blue tit *Cyanistes caeruleus*. To provide a comparable objective measurement of perceived eggshell colours we used an ideal illuminant with equal irradiance across all wavelengths for these analyses. Then we constructed an opponent colour space, using these quantum catches to calculate responses to opponent channels corresponding with relatively short and long wavelength light^35^. Specifically, we calculated the coordinates of eggshell colours within an opponent space defined by perceived eggshell luminance (y axis), and by opponent channels corresponding with the perception of relatively short and long wavelength light^35^ defined by eggshell quantum catches to calculate responses (x axis). Specifically,

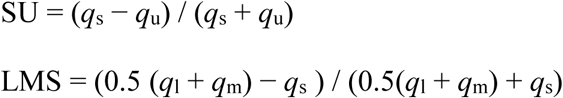

where, *q*_u_, *q*_s_, *q*_m_, *q*_l_ represent the quantile catch of the ultraviolet, short, medium, and long wavelength-sensitive photoreceptors, respectively^35^. The difference between LMS and SU (divided by 2) standardized all colour values to a minimum of −1 and a maximum of +1, corresponding with blue-green and brown respectively (Fig. 1). The eggshell luminance was standardized to the brightest value. Unlike previous analyses that quantified avian eggshell coloration^17,36,37^, this approach uses quantum catch from all four receptors, while providing a quantification of variation in coloration from blue-green to brown, along with a second dimension of capturing variation in perceived eggshell luminance (Fig. 1). Although species differ in their photoreceptor sensitivity, the coordinates within opponent colour spaces were calculated for the average ultraviolet-and violet-sensitive avian viewer, two broadly divergent types of avian vision^38^, were highly correlated (*colour*: β = 0.93, CI_0.95_ = 0.88 to 0.92, *p* < 0.0001, Pagel’s λ _max_=0.85; *luminance*: β = 1.00, CI_0.95_ = 1.00 to 1.00, *p* < 0.0001, Pagel’s λ _max_= 0.85; Extended Data Fig. 2). Our use of avian perceived colour and luminance is important for future studies involving conspecific signalling^21^, brood parasitism^39^, and avian predation^40^. Our calculated avian perceived colour and luminance relate to a measure of blue-green chroma (the sum of the reflectance within 450 and 550nm divided by the sum total reflectance within 300 and 700nm) and brightness (the average reflectance within 300 to 700nm) ^41^ (*colour*: β = −0.97, CI_0.95_ = −0.99 to −0.95, *p* < 0.0001; *luminance*: β = 0.98, CI_0.95_ = 0.98 to 0.99, *p* < 0.0001), respectively. We chose to quantify colour using an avian perceptual system because birds actively select eggs, and therefore these data have value for meta-replication when testing hypotheses related to avian signalling.

### Biogeographical sampling

We downloaded bird distribution maps from Birdlife^42^, and buffered entirely oceanic ranges by 10 km using the ‘Digital Chart of the World’ base map^43^ to restrict all ranges to land. We chose the 10 km buffer because it was well within one sampling area (see below). Then, we removed all lakes from all sampled species bird ranges using a 1:10,000,000 map of worldwide lakes from Natural Earth Data version 3.0.0^44^. We obtained environmental data from WorldClim^45^, National Center for Atmospheric Research^25^ and the Atlas of the Biosphere^46^ datasets. Next we randomly generated one nest every 10,000 km^2^ within each species’ breeding or resident range^42^. Each point was then assigned its species’ colour and luminance value. We overlaid an equal area hexagonal discrete global grid (ISEA aperture 3, resolution 7, *n =* 7,158), and within each hexagon (hereafter locales) we averaged both egg and environmental data. Köppen climate regions^47^ were summarized to each locales’ modal primary category (tropical, arid, temperate, cold, and polar). We pooled cold and polar regions to represent ‘cool’ regions, and pooled the remaining regions as ‘other’. All biogeographic sampling was conducted using ArcGIS ArcMap version 10.5 (Esri, Redlands, CA).

### Phylogenetic and geospatial analysis

We accounted for the phylogenetic relatedness among birds by constructing a phylogenetic hypothesis using a sample of 9,999 fully resolved phylogenetic trees from a recent complete avian phylogeny^48,49^. Using these data, we calculated the Bayesian maximum credibility tree (Extended Data Fig. 3), using the mean branch lengths of the candidate set using DendroPy^50^, dropping 10% of trees as burn in. We then assigned all nodes an ancestral nest type, assuming equal rates using a maximum likelihood estimation^51^. For each locale, we calculated the phylogenetic mean colour and luminance for all birds, as well as birds nesting exclusively on the ground, in open nests, dome nests, cavities, or in mounds using the ‘phyloMean’ function in the ‘motmot 2.0’ package. We removed any locale that could not be phylogenetically controlled for (e.g., fewer than 3 species), which reduced our final sample size to 6,692 locales. Mean species richness across each climate region in our final dataset (*n* = 6,692) were all greater than 15 (*tropical*: 30.50 ± 0.37, *arid*: 24.75 ± 0.42, *temperate*: 36.22 ± 0.65, *cold*: 40.28 ± 0.47, *polar*: 15.32 ± 0.41). Although phylogenetic signal^52^ varied across geography (*colour*: Moran’s I = 0.84, *p* < 0.0001, *luminance*: Moran’s I = 0.89, *p* < 0.0001), phylogenetic means and non-phylogenetic means were highly correlated (*colour*: R^2^ = 0.90, z = 70.43, *p* < 0.0001; *luminance*: R^2^ = 0.91, z = 80.43, *p* < 0.0001; Extended Data Fig. 2a) suggesting that any bias introduced by controlling for full phylogenetic signal (Brownian motion) was minimal. Significance testing for Moran’s I was two tailed.

### Thermoregulation and colour

We tested whether the thermoregulatory properties of the egg could be predicted by eggshell colour and luminance using domestic chicken *Gallus gallus domesticus* eggs (Carrol’s, Pete and Jerry’s, and Stop and Shop) under natural illumination conditions. These eggs (*n =* 48) were either dark brown, light brown, blue-green, or white. Each egg’s mass was recorded using a microbalance (Ohaus Adventure Pro, model AV114C, ± 0.0001g) to the nearest centigram, their surface reflectance was recorded using an Ocean Optics Jaz spectrophotometer (Ocean Optics, Jaz, Dunedin, Florida, USA), and then each egg’s coordinates were calculated within the opponent space. The eggs were then sorted into 12 groups of four, each containing an egg of each colour in a random order. These eggs were acclimated to room temperature ~24.1 °C overnight, and then placed in direct sunlight on 24 August 2018 at 27°C. We measured temperature using a thermal imaging camera (FLIR Infra-Cam) at 15-minute intervals for 75 minutes spanning solar noon (30 min prior to 45 minutes after solar noon). Mean egg temperature for each egg was calculated using calibrated thermal images in ImageJ^53^. To verify that estimates of surface temperature measured by the FLIR Infra-Cam correspond with internal temperatures, we inserted a Omega type T thermocouple (Omega SSRTC-TT-T-24-36) 2.5 cm into a new set of dark brown (*n =* 3) and white eggs (*n =* 3) and recorded internal temperature using a thermocouple logger (Omega HH506RA); internal and external temperature were highly related (r = 0.92, CI_0.95_ = 0.83 to 0.95, *p* < 0.0001; Extended Data Fig. 1*b*).

### Statistical analysis

We accounted for spatial autocorrelation in both dependent and independent variables using a spatial Durbin error model, using first and second order Queen’s contiguity^54^weighting and lower-upper matrix decomposition. A first order Queen’s contiguity considers all neighbours for each locale, while in this case a second contiguity considers all neighbouring locales as well as all their neighbours (weights were roughly equivalent to ~350 km). We use AICc^55,56^ to determine whether the model weighted by the first or second order contiguity better explained our data, and we report the model with the lowest AICc. We calculate and report total impact statistics (e.g., z scores and two tailed p values) and Nagelkerke pseudo-R^2^ for each spatial Durbin error model. All isolated locales and locales with missing information were removed from weight files, which is a requirement of the spatial Durbin error model. This resulted in datasets of eggshell coloration and luminance of the eggs from locales containing all birds (*n =* 6,692), ground nesting birds (*n =* 6,539), open nesting birds (*n =* 6,475), and cavity nesting birds (*n =* 6,557). Phylogenetic mean colour and luminance for each nest type was considered in the presence of all possible nest types at each locale, rather than truncating the dataset. Dome nesting birds and mound builders were retained for these calculations (Extended Data Fig. 3), but we do not have predictions for these groups so we do not explore their independent relationships. We predicted the rate of heating of chicken eggs under natural incident solar radiation based on colour, luminance, and mass using a general linear model.

## Data availability

The data and codes that support the findings of this study are available from the corresponding author upon request.

---

**Extended Data Fig. 1.**
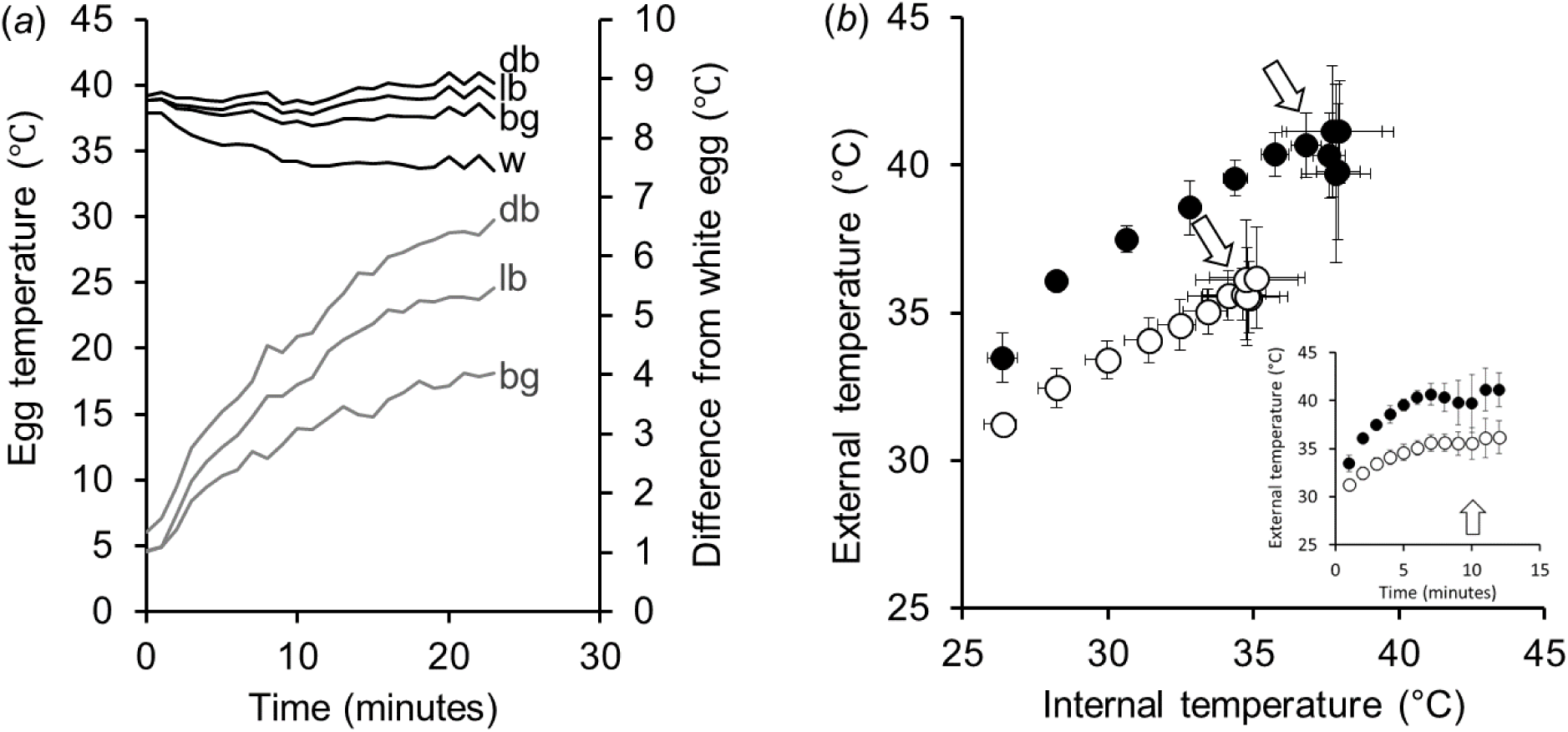
Egg temperatures. Eggshell surface temperatures under **a**, natural ambient light conditions (24.1 °C) for dark brown (db), light brown (lb), blue-green (bg), and white (w) domestic chicken eggs (*n* = 12). All eggs were heated together overnight to 37°C in a Powers Scientific Inc. (model DROS33SD) incubator on locally sourced topsoil, sand, and fallen leaves to approximate a scratch nest. Over this trial, brown eggs retained their intial temperature, while blue-green eggs lost temperature slowly, and white eggs lost temperature rapidly. We measured egg temperatures every minute for 24 minutes using a FLIR Infra-Cam. After 24 minutes, dark brown eggs were 7°C warmer than white eggs (grey lines, second y axis). A new set of eggs’ external surface tmperatures were **b**, related to their internal temperatures. Dark brown (filled dots, *n* = 3) and white (open dots, *n* = 3) eggs were left under natural ambient light conditions (33.9 °C; see Methods for details). The external temperatures change more rapidly than internal temperatures when wind increases convective cooling (arrows), as experienced 10 minutes into this trial (inset).

**Extended Data Fig. 2.**
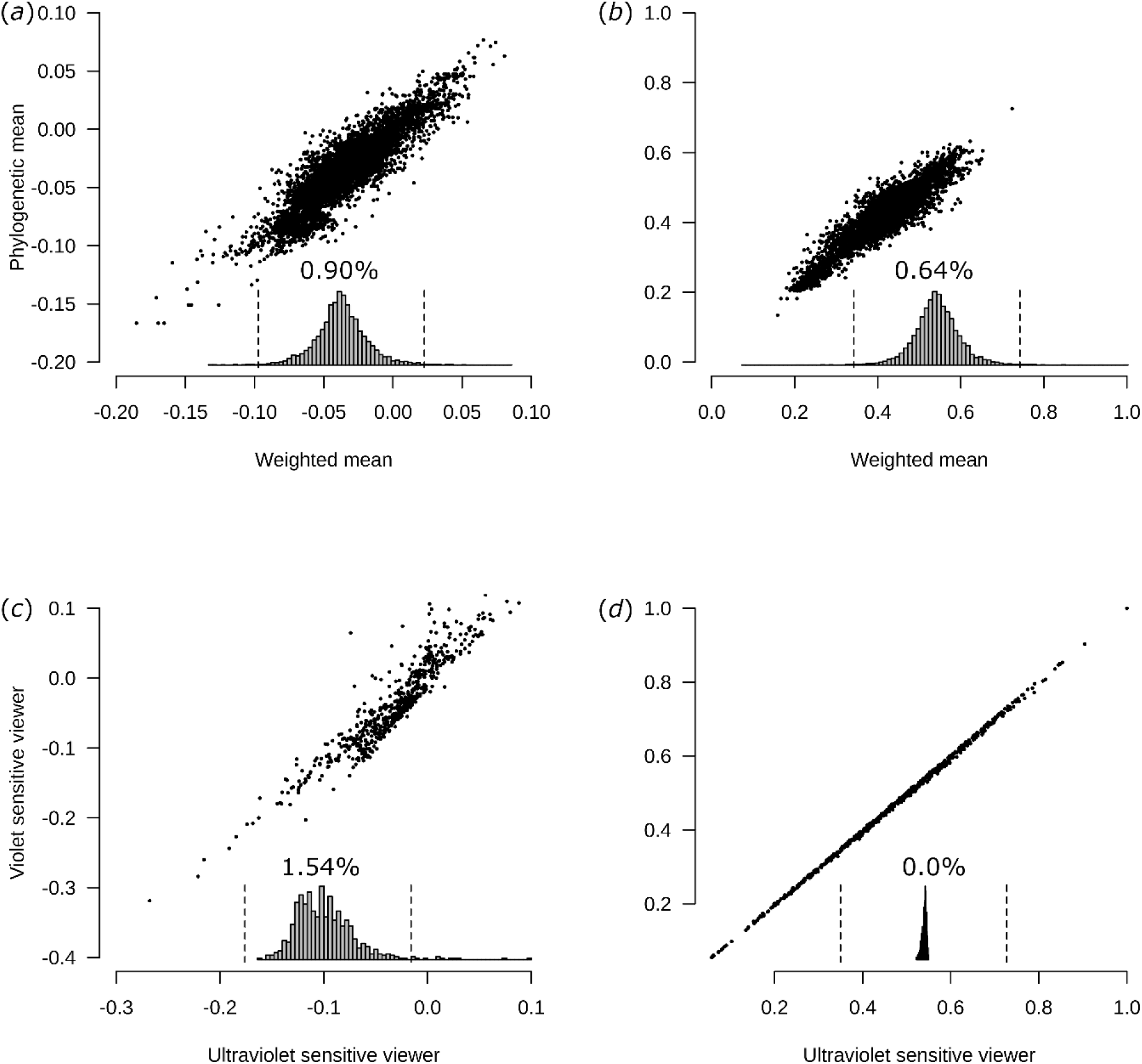
Comparison of other methods. Relationships between the phylogenetic and weighted means for **a**, colour and **b**, luminance, as well as the relationship between the **c**, colour and **d**, luminance components of the opponent colour space calculated for the average violet-and ultraviolet-sensitive avian viewer. Here the weighted means were calculated using an intercept-only phylogenetic generalized least squares, where estimates correspond with weighted means^57^but the maximum likelihood value for Pagel’s lambda^52^is calculated for each locale. These figures illustrate the maximum degree of error introduced into our analyses (residuals) by our application of phylogenetic means and ultraviolet-sensitive visual systems. In both cases, more fine-tuned variation likely exists in our data, but these illustrate extremes (e.g., no phylogenetic correction versus full Brownian motion, and two common but broadly divergent visual systems). The inset histograms represent the residuals from each’s respective tests (see Results). Units are scaled, and presented on a comparable scale (−2.5 to 2.5 standard deviations) and dashed lines represent −1 and 1 standard deviation, respectively. The percentages above each histogram represent deviations more extreme than 1 standard deviation.

**Extended Data Fig. 3.**
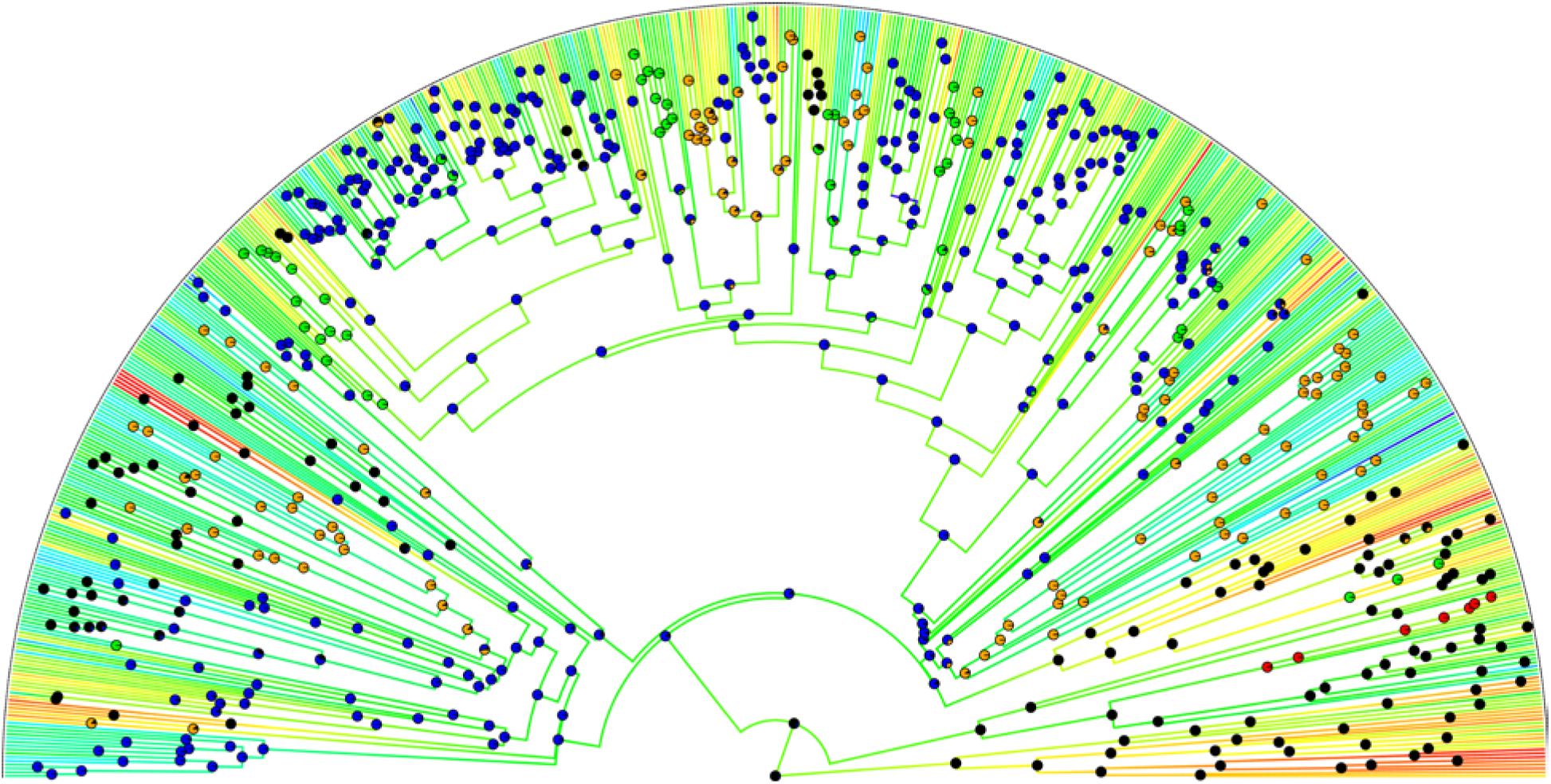
Phylogenetic relationships. The maximum clade credibility tree used in this study. Here we plot branch lengths in continuous coloration representing avian perceived eggshell luminance from dark (red) to bright (blue). At each node, we illustrate pie charts representing the most likely ancestral state for nest types: ground nesting (black), cup nesting (blue), dome nesting (green), cavity nesting (orange), and mound nesting (red).

